# Disease implications of animal social network structure: a synthesis across social systems

**DOI:** 10.1101/106633

**Authors:** Pratha Sah, Janet Mann, Shweta Bansal

## Abstract

1. The disease costs of sociality have largely been understood through the link between group size and transmission. However, infectious disease spread is driven primarily by the social organization of interactions in a group and not its size.
2. We used statistical models to review the social network organization of 47 species, including mammals, birds, reptiles, fish and insects by categorizing each species into one of three social systems, *relatively solitary*, *gregarious* and *socially hierarchical*. Additionally, using computational experiments of infection spread, we determined the disease costs of each social system.
3. We find that relatively solitary species have large variation in number of social partners, that socially hierarchical species are the least clustered in their interactions, and that social networks of gregarious species tend to be the most fragmented. However, these structural differences are primarily driven by weak connections, which suggests that different social systems have evolved unique strategies to organize weak ties.
4. Our synthetic disease experiments reveal that social network organization can mitigate the disease costs of group living for socially hierarchical species when the pathogen is highly transmissible. In contrast, highly transmissible pathogens cause frequent and prolonged epidemic outbreaks in gregarious species.
5. We evaluate the implications of network organization across social systems despite methodological challenges, and our findings offer new perspective on the debate about the disease costs of group living. Additionally, our study demonstrates the potential of meta-analytic methods in social network analysis to test ecological and evolutionary hypotheses on cooperation, group living, communication, and resilience to extrinsic pressures.

## Introduction

Host social behaviour plays an important role in the spread of infectious diseases. Socially complex species from honeybees to African elephants live in large groups and are considered to have elevated costs of pathogen transmission due to high contact rates (Loehle, 1995; Altizer *et al.*, 2003). Previous studies have tested hypotheses about the disease costs of sociality by associating group size with infection transmission (Rifkin, Nunn & Garamszegi, 2012; Patterson & Ruckstuhl, 2013). Beyond a simple dependence on group size, however, recent work in the field of network epidemiology has shown that infectious disease spread largely depends on the organization of infection-spreading interactions between individuals (Godfrey *et al.*, 2009; White, Forester & Craft, 2015; Craft, 2015; VanderWaal & Ezenwa, 2016). Indeed, even when interactions between individuals are assumed to be homogeneous, the expectation of higher disease costs of group living has been mixed (Arnold & Anja, 1993; Rifkin, Nunn & Garamszegi, 2012; Patterson & Ruckstuhl, 2013).

Mathematically, social networks describe patterns of social connections between a set of individuals by representing individuals as nodes and interactions as edges (Croft, James & Krause, 2008; Krause *et al.*, 2014; Farine & Whitehead, 2015). The advantage of social network analysis is that it integrates heterogeneity in interaction patterns at individual, local and population scales to model global level processes, including the spread of social information and infectious diseases (Krause, Croft & James, 2007; Krause *et al.*, 2014; Silk *et al.*, 2017a,b). In recent years, network analysis tools have allowed for rapid advances in our understanding of how individual interaction rates are related to the risk of acquiring infection (Otterstatter & Thomson, 2007; Leu, Kappeler & Bull, 2010). A fundamental individual-level characteristic relevant to the spread of social or biological conta-gion in networks is the number of direct social partners, associates or contacts, capturing the interaction necessary for transmission. While much attention has been focused on the implications of individual sociality, the disease implications of a species’ social system remains unclear.

By quantifying group-level metrics that describe global structures in interaction patterns, the network approach provides a unique opportunity to examine the disease costs of species social system. The role of higher-order network structures such as degree heterogeneity (Fig. 1A), subgroup cohesion (Fig. 1D), network fragmentation (Fig. 1E), and average clustering coefficient (Fig. 1F) on infectious disease spread is complex, but is relatively well understood (see network structure definitions in Table S1)(Keeling, 2005; Meyers *et al.*, 2005; Sah *et al.*, 2017). For example, as degree heterogeneity (or variation in the number of social partners) in a network increases, the epidemic threshold (i.e., the minimum pathogen trans-missibility that can cause large outbreaks) decreases (Anderson, May & Anderson, 1992). However, the probability of epidemic outbreaks is lower in networks with high degree variance for moderately and highly transmissible pathogens (Meyers *et al.*, 2005). Network metrics such as average clustering coefficient, subgroup cohesion and network fragmentation capture the tendency of individuals to form cliques and subgroups (Fig. 1). Although the dynamics of infectious disease spread remain largely unaffected in networks with moderate levels of clustering, cohesion and fragmentation, extreme levels of these metrics in networks reduce epidemic size and prolong epidemic outbreaks (Keeling, 2005; Sah *et al.*, 2017).

**Fig. 1.**
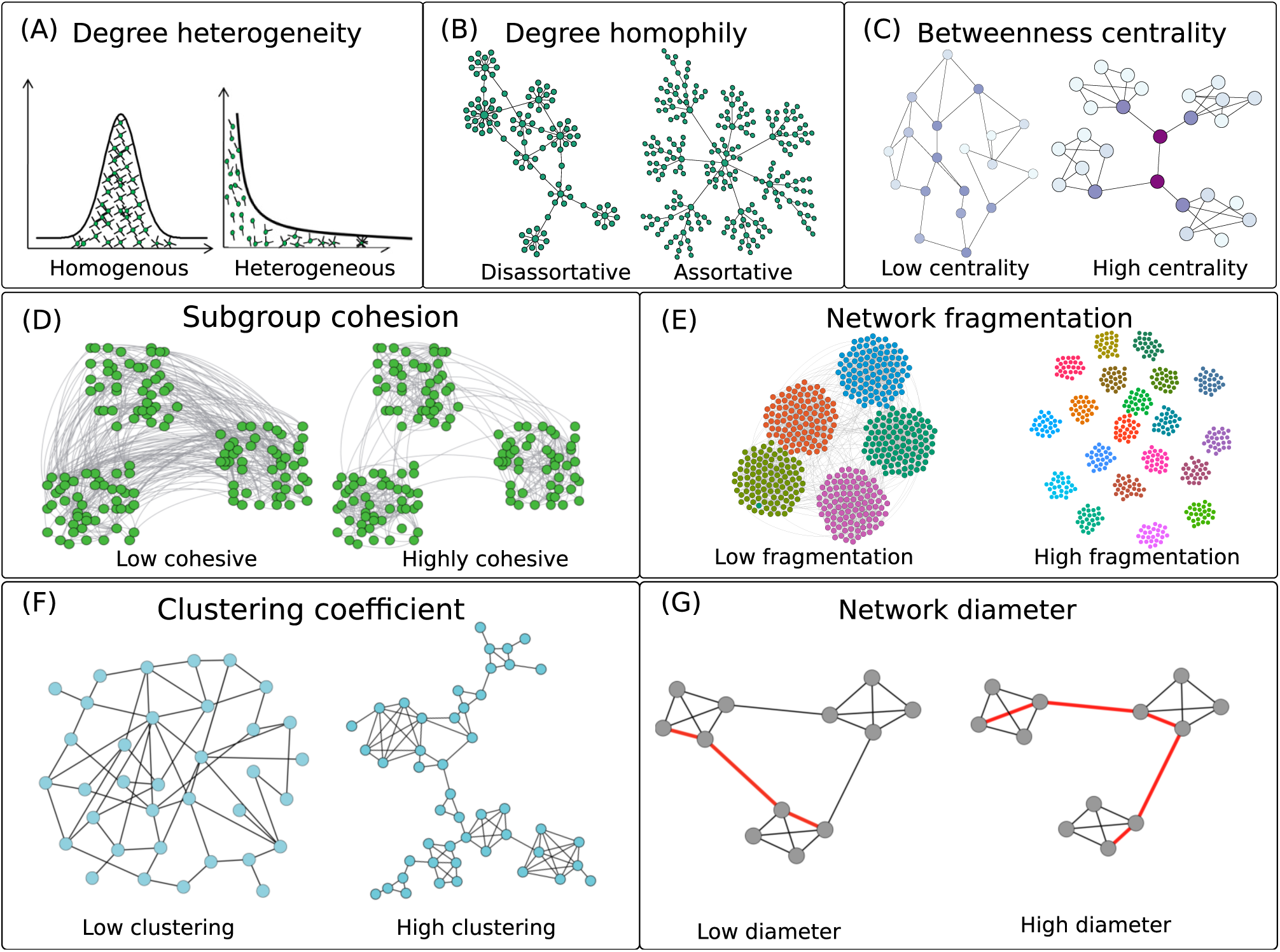
A stylized illustration of the global network measures used (in the final model) to identify the structural differences in the social networks among different social systems. (A) Degree heterogeneity, measured as the coefficient of variation (CV) in the frequency distribution of the number of social partners (known as the *degree distribution*). Shown is the degree distribution of a homogeneous network (CV ≪ 1), and an exponential degree distribution of a network with large varia-tion in individual degrees (CV = 1). (B) Degree homophily (*ρ*), or the tendency of social partners to have a similar degree. Shown is an example of a disassortative network, wherein high degree individuals tend to associate with low degree individuals (*ρ* < 0), and assortative degree networks, where high degree individuals tend to form social bonds with each other (*ρ* > 0). (C) Average betweenness centrality, that measures the tendency of nodes to occupy central position within the social network. Shown is an example of a network with low average betweenness centrality and a network with high average betweenness centrality. Node colors represent the betweenness centrality values - nodes with darker colors occupy more central positions within the network. (D) Subgroup cohesion measures the tendency of individuals to interact with members of own subgroups (modules). The network to the left has three low cohesive subgroups, while the network to the right has highly cohesive subgroups where most of the interactions occur within (rather than between) subgroups. (E) Network fragmentation, measured as the log-number of the subgroups (modules) present within the largest connected component of a social network. Shown is an example of low (left) and highly (right) fragmented network. (F) The average clustering coefficient measures the average fraction of all possible triangles through nodes that exist in the network, and indicates the propensity of social partners of individuals to interact with each other. (G) Network diameter is the longest of all shortest paths between pairs of nodes in a network. Shown is an example of a network with low network diameter (longest of shortest paths = 3) and a similar network with network diameter of 5, indicated by red coloured edges.

Recent mathematical models predict that the network structure of socially complex species can serve as a primary defence mechanism against infectious disease by lowering the risk of disease invasion and spread (Hock & Fefferman, 2012). It remains uncertain, however, whether the structure of social networks naturally observed in less-complex social systems mediates infectious disease risk and transmission. A systematic examination of the disease costs associated with species social system requires a comparative approach that isolates unique structural characteristics of social connections, while controlling for population size, data collection methodology and type of interaction recorded. However, comparing networks across different taxonomic groups has proven to be a difficult task, with only a few cross-species network comparisons previously published in the literature (Faust & Skvoretz, 2002; Faust, 2006; Sah *et al.*, 2017).

In this study, we conduct a quantitative comparative analysis across 47 species to investigate whether social network organization alone, without the presence of physiological or behavioural immune responses, can reduce the disease costs of group living for various social systems. This is achieved in three steps. First, we categorize the continuum of species sociality into three distinct social systems (relatively solitary, gregarious and socially hierarchical); we then use phylogenetically-controlled Bayesian generalized linear mixed models to identify social network structures which are predictive of the three social systems. Second, we perform computational experiments of infection spread to compare epidemiological outcomes (epidemic probability, epidemic duration and epidemic size) associated with the identified social network structures. In the final step, we investigate whether the differences in these network structures across the three social systems trans-lates to differences in their disease outcomes.

We hypothesize that a social species can mitigate disease costs associated with group living through the organization of their social structure. However, we expect the presence of alternate disease defence mechanisms to also play an important role: social insects, for example, use social immunity as a primary strategy to minimize disease transmission; the structure of the social network in such species may not be effective in preventing future outbreaks or reducing disease transmission. Our analysis, by broadening the scope of network analysis from species-specific analysis to a meta-analytic approach, offers new perspective on how social structure strategies mediate the disease costs of group living. A better understanding of the association between network structure and different social systems can facilitate investigations on other evolutionary and ecological hypotheses on group living, social complexity, communication, population robustness and resilience to extrinsic population stressors.

## Materials and methods

### Dataset

We first conducted electronic searches in *Google Scholar* and popular data repositories, including *Dryad Digital Repository* and *figshare* for relevant network datasets associated with peer-reviewed publications. We used the following terms to perform our search: "social network", "social structure", "contact network", "interaction network", "network behaviour", "animal network", "behaviour heterogeneity" and "social organization". Only studies on non-human species were considered in our primary search. Network studies not reporting interactions (such as biological networks, food-web networks) were excluded. By reviewing the quality (i.e., whether enough information was provided to accurately reconstruct networks) of published networks datasets, we selected 666 social networks spanning 47 animal species and 18 taxonomic orders. Edge connections in these networks represented several types of interactions between individuals, including dominance, grooming, physical contact, spatial proximity, direct food-sharing (i.e. trophallaxis), foraging, and interactions based on the asynchronous use of a shared resource. Fig. 2 summarizes the species, the number of networks and the reported interaction types contributed by each taxonomic order represented in the study.

**Fig. 2.**
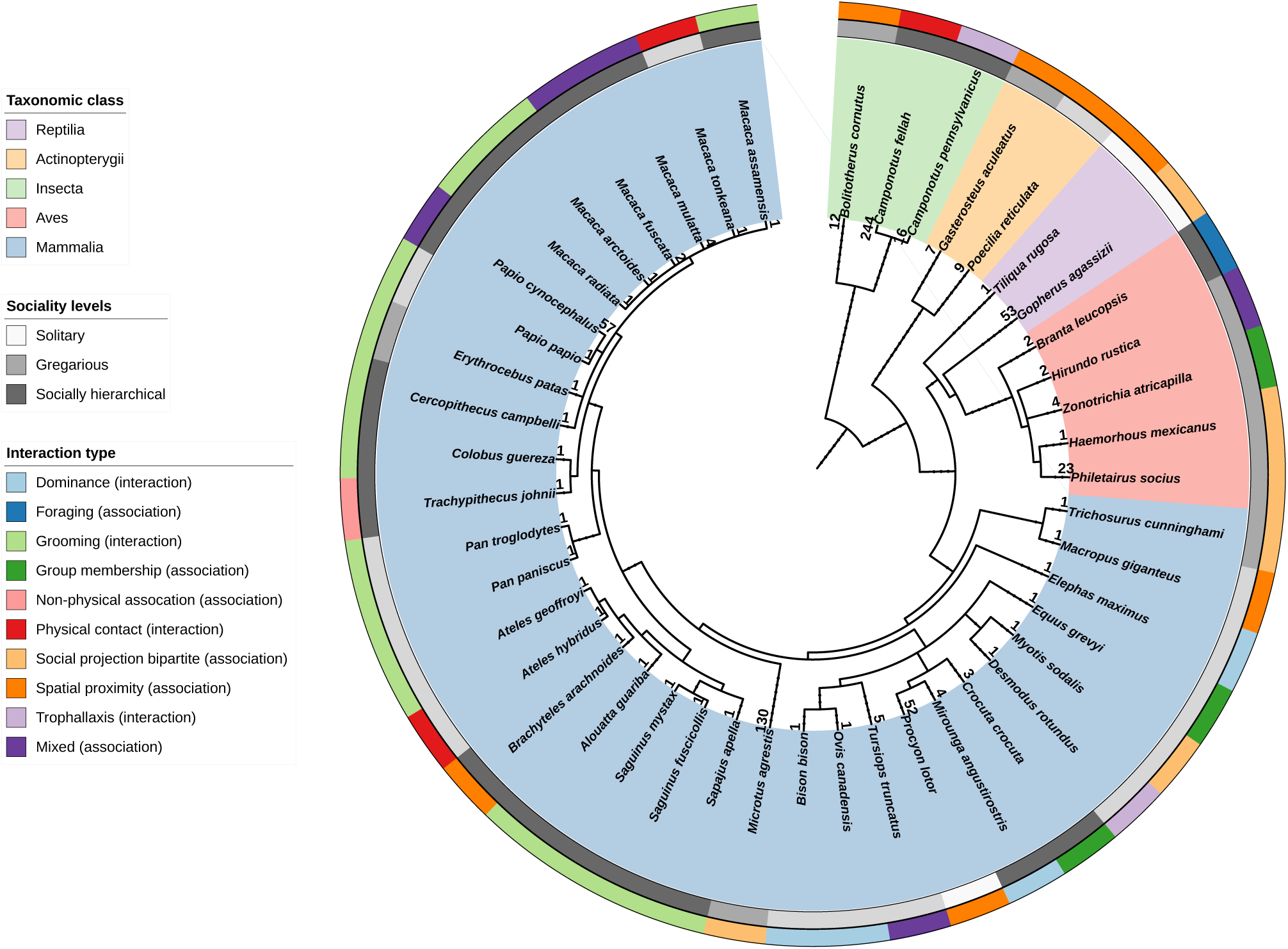
Phylogenetic distribution of animal species represented in the social network dataset used in this study. Numbers next to the inner ring denote the total networks available for the particular species. The inner and the middle ring is color coded according to the taxonomic class and the social system of the species. The colors in the outer ring indicates the type of interaction represented in the network, and whether the interactions were coded as (direct) interactions or association in our analyses (in brackets). The tree was constructed in the In-teractive Tree Of Life (http://itol.embl.de/) from the NCBI taxonomy database (http://www.ncbi.nlm.nih.gov/Taxonomy/).

### Classifying species’ social system

Developing a definition of social structure that encompasses the continuum of social systems across diverse taxonomic groups is challenging. Consequently, we followed Slater & Halliday (1994) and Kappeler & van Schaik (2002) to classify species into three broad categories of social structure based on the degree of association between adults during activities such as foraging, travelling, sleeping/resting and rearing offspring. *Relatively solitary* species were defined by infrequent aggregation or association between adults outside of the breeding period, and lack of synchronized movements in space by adults. Examples of relatively solitary species in the database include the desert tortoise (*Gopherus agassizii*), wild raccoons (*Procyon lotor*), and the Australian sleepy lizard (*Tiliqua rugosa*). Recent studies suggest that the social structure of a species traditionally considered as solitary can be complex (Sah *et al.*, 2016; Prange *et al.*, 2011). We therefore categorized the three species as *relatively solitary* and not solitary. Species that aggregate for one or more activities, but have unstable or temporally varying group composition were classified as *gregarious*. Examples of gregarious species in our database include bottlenose dolphins (*Tursiops truncatus*), bison (*Bison bison*), Indiana bats (*Myotis sodalis*), female Asian elephants (*Elephas maximus*), sociable weavers (*Philetairus socius*), golden-crowned sparrows (*Zonotrichia atricapilla*) and guppies (*Poecilia reticulata*). Species characterized by a permanent or long-term (i.e., at least over a single breeding season) stable social hierarchy were classified as *socially hierarchical*. Examples of socially hierarchical species include carpenter ants (*Camponotus fellah*), yellow baboons (*Papio cynocephalus*), male elephant seals (*Mirounga angustirostris*) and spotted hyenas (*Crocuta crocuta*). We note that animal social behaviour is being increasingly recognized to span a continuum from solitary to eusocial (Aureli *et al.*, 2008; Aviles & Harwood, 2012; Silk, Cheney & Seyfarth, 2013), with most species showing some level of fission-fusion dynamics (Silk *et al.*, 2014). The division of social systems into three discrete, albeit arbitrary, categories allows for simple distinctions in the organization of network structure and disease risks among species that are characterized by different complexity in group living behavior.

### Identifying unique network structures of species’ social system

To examine the structure of social networks associated with our three classified social systems, we used a Bayesian generalized linear mixed model (GLMM) approach using the *MCMCglmm* package in *R* (Hadfield, 2010), with the species’ social system as the response (categorical response with three levels - relatively solitary, gregarious and socially hierarchical). The following network measures were included as predictors in the model (see Table S1 in Supporting information for definitions and Fig.1 for illustrations): degree heterogeneity, degree homophily, average clustering coefficient, weighted clustering coefficient, transitivity, average betweenness centrality, weighted betweenness centrality, average subgroup size, network fragmentation, subgroup cohesion, relative modularity and network diameter. Network fragmentation (i.e., the number of subgroups within the largest connected component of the social network) and Newman modularity was estimated using the Louvain method (Blondel *et al.*, 2008). Relative modularity was then calculated by normalizing Newman modularity with the maximum modularity that can be realized in the given social network (Sah *et al.*, 2014, 2017). The rest of the network metrics were computed using the *Networkx* package in Python (https://networkx.github.io/). We controlled for network size and density by including the number of nodes and edges as predictors, and mean edge weight was included to control for data sampling design. To control for phylogenetic relationships between species, a correlation matrix derived from a phylogeny was included as a random factor. The phylogenetic relationship between species was estimated based on NCBI taxonomy using phyloT (http://phylot.biobyte.de). We controlled for repeated measurements within groups, animal species, the type of interaction recorded, and edge weighting criteria by including *group*, *taxa*, *interaction type* (association *vs.* interaction) and *edge weight type* (weighted *vs.* unweighted) as random effects in the analysis. As the spatial scale of data collection can influence network structure (Table S3, Supporting information), we specified sampling scale (social sampling *vs.* spatial sampling) as random effect in all our analyses. Studies that collected data on specific social groups were categorized as *social sampling*, and those that sampled all animals within a fixed spatial boundary were labelled as *spatial sampling*.

All continuous fixed-effects were centered (by subtracting their averages) and scaled to unit variances (by dividing by their standard deviation) to assign each continuous predictor with the same prior importance in the analysis (Schielzeth, 2010). Since network measures can be highly correlated to each other, variance inflation factor (VIF) was estimated for each covariate in the fitted model, and covariates with VIF greater than 5 were removed to avoid multicollinearity. We used a weakly informative Gelman prior for fixed effects and parameter-expanded priors for the random effects to improve mixing and decrease the autocorrelation among iterations (Gelman, 2006). Specifically, a *χ*^2^ distribution with 1 degree of freedom was used as suggested by Hadfield (2014). We ran three MCMC chains for 15 million iterations, with a thinning interval of 1000 after burn in of 50,000. Convergence of chains was assessed using the Gelman-Rubin diagnostic statistic (Gelman & Rubin, 1992) in the *coda* package (Plummer *et al.*, 2006).

Groups of certain species in our database were represented with multiple networks, each summarizing a set of interactions occurring in a discrete time period. To ensure that such animal groups were not over-represented in the original analysis, we performed a cross-validation of our analysis by random sub-sampling. Specifically, we repeated the analysis 100 times with a random subset of the data composed of (randomly selected) single networks of each unique animal group in our database. An average of coefficient estimates across the multiple subsam-ples was then calculated and compared to the coefficients estimated using the full dataset.

### Evaluating the role of weak ties in driving structural differences in species’ social system

The analysis described in the previous section assumes equal importance of all edges recorded in a social network. To examine the role of weak ties in driving the structural differences between the three social systems, we removed edges with weights lower than a specified threshold. Four edge weight thresholds were examined in detail: 5%, 10%, 15% and 20%. Specifically, all edges with weights below the specified threshold were removed to obtain thresholded social networks. For example, to construct a 10% threshold network from an original network with maximum edge weight ω, we removed all edges with weights below 0.1 × *ω*. Next, the phylogenetically-controlled Bayesian mixed model analysis described in the previous section was repeated to determine the structural difference between the thresholded networks of the three social systems. We ran four separate models, each with one of the four thresholds.

### Disease implications of network structure and species’ social system

We considered disease costs of the three social systems with synthetic experiments based on a computational disease model, and followed up with statistical analysis of the results.

### Disease simulations

We performed Monte-Carlo simulations of a discrete-time susceptible-infected-recovered (SIR) model of infection spread through each network in our database. For disease simulations, we ignored the weights assigned to social interactions between individuals, because the impact of interaction weight (whether they represent contact duration, frequency or intensity) on infection spread is generally not well understood epidemiologically. Transmissibility of the simulated pathogen was defined as the probability of infection transmission from an infected to susceptible host during the infectious period of the host. Assuming infection transmission to be a Poisson process and a constant recovery probability (Grenfell & Dobson, 1995; Kiss, Miller & Simon, 2017), the pathogen transmissibility can be calculated as 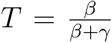, where *β* and *γ* is the infection and recovery probability parameter, respectively (Bansal, Grenfell & Meyers, 2007). The stochastic epidemiological simulations used in this study are based on a discrete-time, chain binomial, SIR model (Bailey, 1957). Each disease simulation was initiated by infecting a randomly chosen individual in the social network. At subsequent time steps every infected individual in the network could either transmit infection to a susceptible neighbour with probability parameter *β* or recover with probability *γ*. The disease simulations were terminated when there were no remaining infected individuals in the network. We performed disease simulations with a wide range of transmissibility values (0.05 to 0.45, with increments of 0.05), by varying infection probability (*β*) and assuming a constant recovery probability (*γ* = 0.2 or average infectious period of 5 days). In the paper, we focus our discussion on three specific values of pathogen transmissibility (*T* = 0.05, 0.15, and 0.45) because they correspond to low, moderate and highly contagious infectious diseases with average basic repro-duction numbers (R0) of 1.6, 4.6 and 14.0, respectively (Heffernan, Smith & Wahl, 2005). The detailed results of disease simulations over a wider range of pathogen transmissibility (0.05 – 0.45) are included in the Supporting information.

To investigate the effects of recovery probability on the behavior of pathogen spread, we repeated disease simulations with a similar range of transmissibility values as before (0.05 to 0.45), but with a longer infectious period (10 days or *γ* = 0.1). For each combination of pathogen transmissibility and social network, 500 simulations of disease spread were carried out and summarized using three measures: (a) epidemic probability, the likelihood of an infectious disease invasion turning into a large epidemic (outbreaks that infect at least 15% of the population) (b) epidemic duration, the time to epidemic extinction, and (c) epidemic size, the average percentage of individuals infected in an epidemic outbreak.

### Evaluating disease outcomes of network structure and species’ social system

Three separate linear Gaussian models, one corresponding to each outbreak measure (epidemic probability, epidemic duration, and epidemic size), were fit to establish disease costs of network measures associated with species’ social system using using the R package MCMCglmm (Hadfield, 2010). To evaluate the role of network structure on the probability of large outbreaks, pathogen transmissibility and network measures included in the final model of the previous analysis were included as predictors (Table1). We repeated the analysis with the species’ social system as predictor to directly estimate the vulnerability of different social structure towards disease transmission.

In all models, the effective number of nodes (i.e., the number of individuals with degree greater than zero), network density and the size of the largest connected component of the network were also included as controlling predictors. As before, we controlled for the presence of phylogenetic correlations, group identification, animal species, edge weight type, and sampling scale of networks. As infectious disease spread over different interaction types represents different transmission routes, we also controlled for pathogen transmission mode by including the interaction type as a random effect. Minimally informative priors were used for fixed effects (normal prior) and (co)variance components (inverse Wishart; Had-field (2010)). We ran three MCMC chains for 100 thousand iterations, with a thinning interval of 10 after burn-in of 2000, and assessed convergence using the Gelman-Rubin diagnostic statistic (Gelman & Rubin, 1992) in the *coda* package. To make posthoc comparisons within the models, we performed pairwise comparisons between the three social systems with a Tukey adjustment of *P* values, using the *lsmeans* R package (Lenth, 2016).

## Results

### Unique network structures associated with species’ social system

The final model (after removing collinear predictors) consisted of seven global network measures - degree heterogeneity, degree homophily, average betweenness centrality, average clustering coefficient, subgroup cohesion, network fragmentation and network diameter (Fig. 1, Table 1). Out of the five random effects included in the model (phylogeny, group identification, interaction type, edge type, sampling scale), phylogeny explained a large portion of the variance (Table S2, Supporting information), indicating that there is a substantial phylogenetic correlation within the social systems. Of the three social systems (relatively solitary, gregarious and socially hierarchical), the social networks of relatively solitary species demonstrated the largest variation in the number of social partners, or degree heterogeneity (Table 1). In contrast, socially hierarchical species had the least variation in number of social partners, and experienced a local social environment that is not as well inter-connected; this is evident by the low average clustering coefficient of their social networks as compared to other social systems (average clustering coefficient, Table 1). In terms of network fragmentation (which was calculated on the largest connected component of networks), the social networks of gregarious species were the most subdivided into socially cohesive groups. No statistically significant differences were observed between the social systems with respect to other network metrics. Table S3 of Supporting information reports the average coefficient estimates of all seven global network metrics from the cross-validation analysis; all estimates were within the 95% credible interval of the effect sizes reported in the full model (Table 1). We also find that the organization of social networks depends on the sampling scale of social associations, but not on the type of interactions recorded (including when the interaction types are grouped into two categories of direct interactions vs. associations, and when the recorded interactions are categorized into ten distinct types mentioned in Fig. 2). For example, networks measured at a population scale rather for social groups tended to have low local connectivity, as measured by the average clustering coefficient, and low average betweenness centrality (Table S4, Supporting information).

**Table 1.** Effect size estimates of the Bayesian generalized linear mixed models examining the characteristics of social network structure among the three social systems: relatively solitary, gregarious and socially hierarchical. Shown are the posterior means of the expected change in log-odds of being in focal social system (column headers), as compared to the base social system (row headers), with one-unit increase in the network measure. The 95% credible intervals (i.e., the coefficients have a posterior probability of 0.95 to lie within these intervals) are included in brackets. Significant terms with pMCMC < 0.05 are indicated in bold, where pMCMC is the proportion of MCMC samples that cross zero.

| Degree heterogeneity | Focal | Relatively solitary | Gregarious | Socially hierarchical |
| --- | --- | --- | --- | --- |
|  | Base |  |  |  |
|  | Relatively solitary |  | **-3.96 [-7.57, -0.33]** | **-9.46 [-15.21, -3.87]** |
|  | Gregarious |  |  | **-6.39 [-11.67, -1.34]** |
|  | Socially hierarchical |  |  |  |
| Degree homophily | Focal | Relatively solitary | Gregarious | Socially hierarchical |
|  | Base |  |  |  |
|  | Relatively solitary |  | -0.18 [-1.66, 1.17] | -1.69 [-3.80, 0.25] |
|  | Gregarious |  |  | -1.64 [-3.25, 0.09] |
|  | Socially hierarchical |  |  |  |
| Average betweenness centrality | Focal | Relatively solitary | Gregarious | Socially hierarchical |
|  | Base |  |  |  |
|  | Relatively solitary |  | 0.68 [-2.31, 3.76] | 0.36 [-2.91, 3.82] |
|  | Gregarious |  |  | 0.27 [-2.56, 2.12] |
|  | Socially hierarchical |  |  |  |
| Average clustering coefficient | Focal | Relatively solitary | Gregarious | Socially hierarchical |
|  | Base |  |  |  |
|  | Relatively solitary |  | -0.06 [-2.49, 2.47] | **-3.40 [-6.56, -0.24]** |
|  | Gregarious |  |  | **-3.30 [-5.82, -0.88]** |
|  | Socially hierarchical |  |  |  |
| Subgroup cohesion | Focal | Relatively solitary | Gregarious | Socially hierarchical |
|  | Base |  |  |  |
|  | Relatively solitary |  | -0.60 [-2.98, 1.84] | -0.40 [-3.23, 2.42] |
|  | Gregarious |  |  | 0.97 [-1.14, 3.05] |
|  | Socially hierarchical |  |  |  |
| Network fragmentation | Focal | Relatively solitary | Gregarious | Socially hierarchical |
|  | Base |  |  |  |
|  | Relatively solitary |  | **3.94 [0.74, 7.26]** | 0.11 [-4.01, 4.12] |
|  | Gregarious |  |  | **-3.27 [-6.11, -0.51]** |
|  | Socially hierarchical |  |  |  |
| Network diameter | Focal | Relatively solitary | Gregarious | Socially hierarchical |
|  | Base |  |  |  |
|  | Relatively solitary |  | -1.79 [-5.00, 1.45] | 1.46 [-2.79, 5.52] |
|  | Gregarious |  |  | 2.86 [-0.31, 5.89] |
|  | Socially hierarchical |  |  |  |

### Disease costs of network structure and species’ social system

Our previous analysis revealed that only a few features of social networks are significant in distinguishing the three social systems. Next we ask: Do these key topological differences mediate differential disease costs of each social system? To answer this question, we first examined how degree heterogeneity, clustering coefficient and network fragmentation influence epidemic risk and transmission of low, moderate and highly transmissible pathogens (Fig. 3; see Fig. S2, S4 in Supporting information for results on an extended range of pathogen transmissibility values and Fig. S5 for results on disease simulations with extended infectious period). High variation in individual sociality (i.e., high degree heterogeneity) in social networks was predictive of small and short epidemic outbreaks for low transmissible pathogens. Moderately spreading pathogens in network with high degree heterogeneity led to less frequent, shorter epidemics that infected a smaller proportion of the population (degree heterogeneity, Fig. 3). The presence of cliques in social networks was associated with prolonged but small outbreaks of low transmissible pathogens, and higher epidemic risk of moderately transmissible infections (average clustering coefficient, Fig. 3). Subdivisions of networks into socially cohesive groups (high fragmentation) was associated with reduced risk of lowly transmissible infections becoming large epidemics; outbreaks that did reach epidemic proportion were shorter and infected a lower proportion of the population. Conversely, highly contagious pathogens caused frequent, large, and prolonged epidemic outbreaks in networks with high network fragmentation (network fragmentation, Fig. 3).

**Fig. 3.**
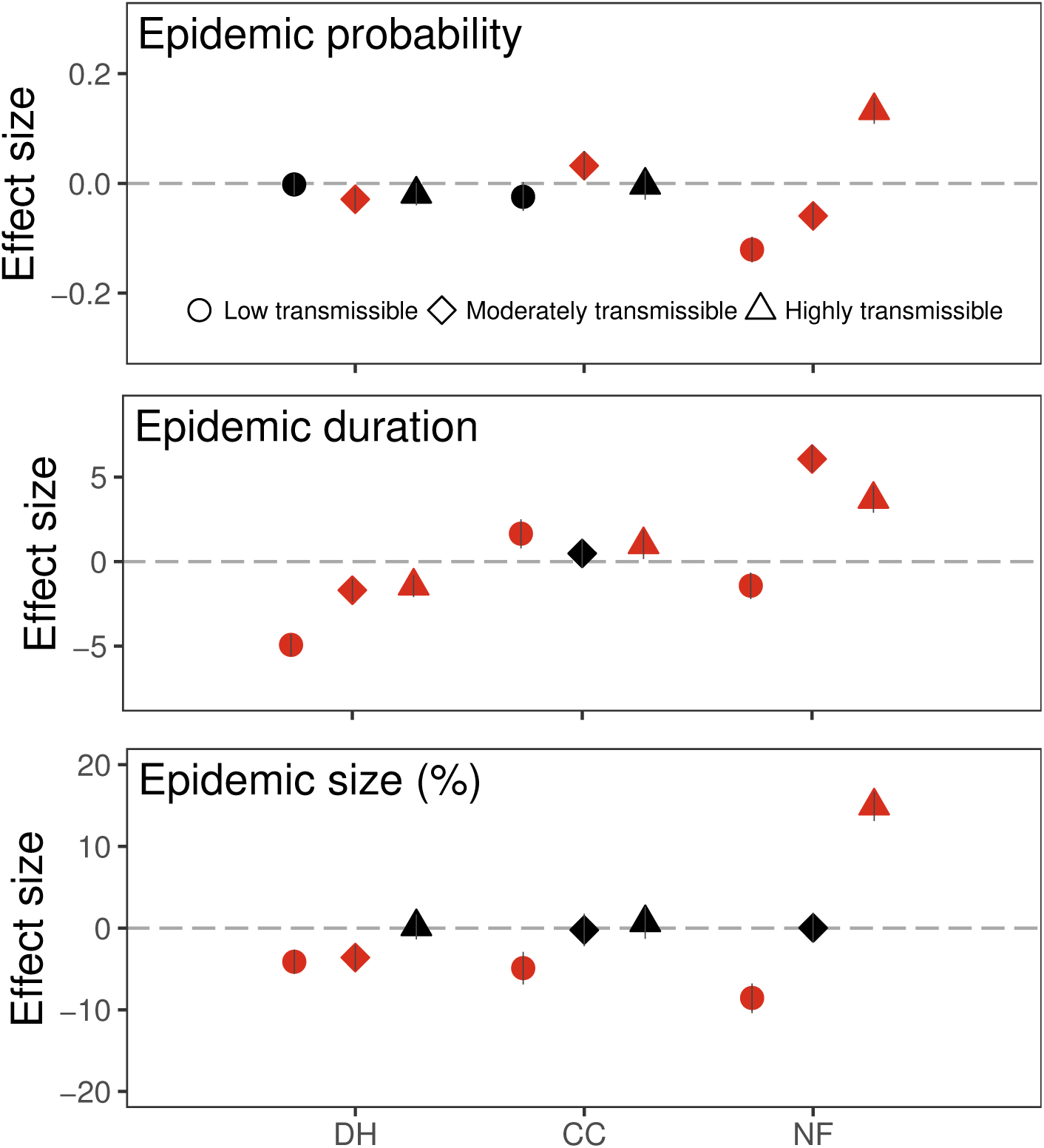
Role of network structures in influencing disease transmission summarized as epidemic probability (likelihood of large outbreaks infecting at least 15% of individuals in the network), average epidemic duration (time to epidemic extinction), and average epidemic size (percent of individuals infected in the social network), for low (=0.05), moderate (=0.15) and highly (=0.45) transmissible pathogens. The average infectious period of the simulated disease is 5 days (*γ*=0.2). The three global network measures shown are the ones that were found to differ among the three social systems (Table 1). DH, degree heterogeneity; CC, average clustering coefficient; NF, network fragmentation. Error bars represent 95% credible intervals. Credible intervals that do not include zero suggest significant association with disease transmission (red = significant effect, black = effect not significant)

Consequently, socially hierarchical species experienced elevated risk of epidemic outbreaks of moderately transmissible pathogen due to homogeneous individual connectivity (low degree heterogeneity) and high global connectivity (low net-work fragmentation) nature of their social networks (epidemic probability, Fig. 4, Fig. S3 and S5 in Supporting information). The highly fragmented networks of gregarious species were more vulnerable to frequent, large, and prolonged epidemic outbreaks of highly transmissible pathogens as compared to other social systems. Given that degree heterogeneity and network fragmentation is associated with shorter outbreaks of low transmissible pathogens (Fig. 3, Fig. S3 and S6 in Supporting information), epidemic duration of less transmissible pathogens was lowest in gregarious species, followed by relatively solitary species (epidemic duration, Fig. 4, Fig. S3 and S6 in Supporting information). For moderately contagious pathogens, highly fragmented networks of gregarious species experienced longer epidemic outbreaks as compared to relatively solitary and socially hierarchical species.

**Fig. 4.**
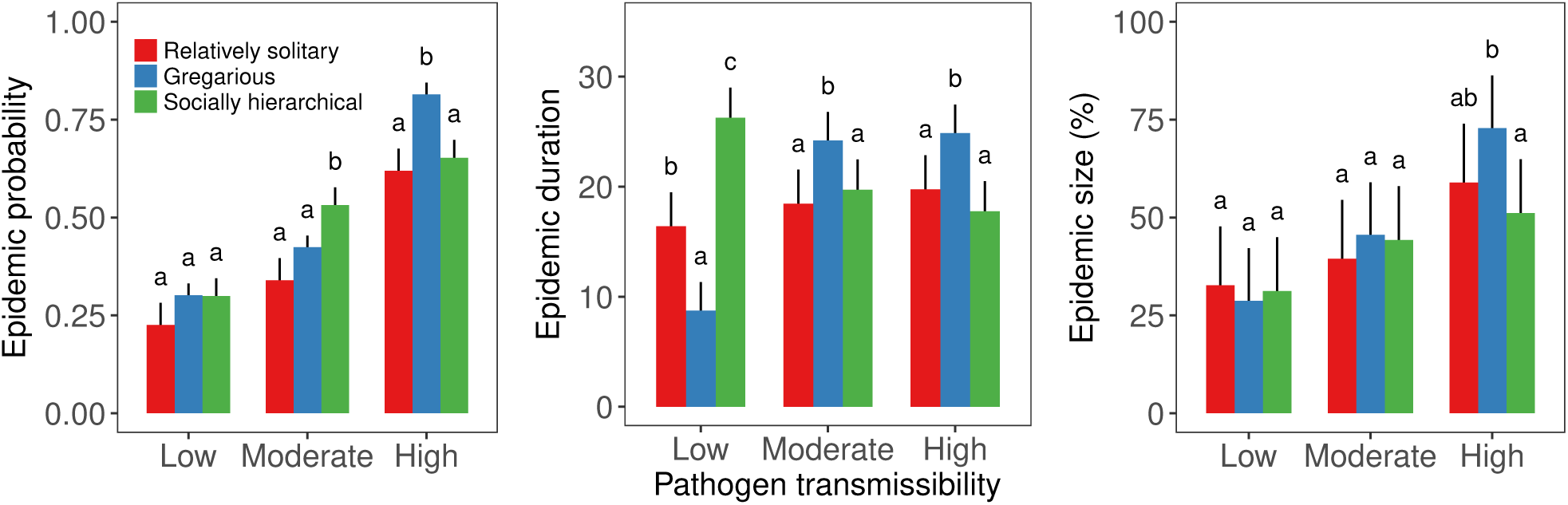
Disease costs of social systems due to social network structure. Disease cost has been quantified in terms of epidemic probability, average epidemic du-ration and average epidemic size for low (=0.05), moderate (=0.15) and highly (=0.45) transmissible pathogens. The average infectious period of the simulated disease is 5 days (*γ*=0.2). Error bars represent standard errors, and different letters above the bars denote a significant difference between the means (P < 0.05)

### Role of weak ties in distinguishing species’ social system, and disease implications

When the weakest 5% edges were removed from all weighted networks, the structural differences between the three social systems were observed mainly in two network metrics - degree heterogeneity and network fragmentation. Similar to the empirical networks (Table 1), the 5% thresholded social networks of relatively solitary species demonstrated the highest variation in number of social partners; and 5% thresholded networks of gregarious species were more fragmented compared to relatively solitary and socially hierarchical species (Table S5, Supporting information). When the weakest 10% and 15% edges were removed, the global network measures across all social systems were similar to each other, except for one important difference. Both 10% and 15% thresholded networks of social species (gregarious and socially hierarchical) demonstrated a statistically significant higher average betweenness centrality, or higher global connectivity than relatively solitary species (Table S6, S7 and S8, Supporting information).

Disease simulations through 20% edge weight thresholded social networks revealed no differences in epidemiological outcomes between the three social systems for all except low pathogen transmissibility (Fig. S7, Supporting information). For slow spreading pathogens, networks of relatively solitary species experienced prolonged epidemic outbreaks as compared to social species.

## Discussion

It is becoming increasingly clear that the impact of an infectious disease on a population depends on the organization of infection-spreading interactions between individuals rather than group size. (Godfrey *et al.*, 2009; Craft, 2015; White, Forester & Craft, 2015; Sah *et al.*, 2017). Since organization of social network structure concurrently impacts the transmission of information and infectious diseases, it has critical implications for understanding the evolutionary tradeoffs between social behavior and disease dynamics. The disease implications of social network structure can differ depending on the evolutionary trajectory of social systems. For instance, social complexity can emerge as a result of selective pressures of past infectious diseases, and therefore may have the ability to lower the risk of transmission of future infectious disease (Hock & Fefferman, 2012). Conversely, the patterns of social interactions may not provide protection from disease transmission in species that use alternate defense mechanisms (physiological or behavioral) to combat disease spread once it is introduced in the population (Cremer, Armitage & Schmid-Hempel, 2007; Stroeymeyt, Casillas-Pérez & Cremer, 2014; Meunier, 2015). In this study, we assessed whether network structure alone (in absence of physiological or behavioral disease defense mechanisms) can reduce the risk of infectious disease transmission in different social systems, using comparative methods on an extensive database of animal social networks.

Our analysis compares global structural features associated with social networks of species classified into three social systems: relatively solitary, gregarious and socially hierarchical. The evidence that we present here suggests that, at the least, relatively solitary, gregarious, and higher social organizations can be distinguished from each other based on *(i)* degree of variation among social partners (i.e. degree heterogeneity), *(ii)* local connectivity, as indicated by the presence of cliques within the social networks (i.e, average clustering coefficient), and *(iii)* the extent to which the social network is divided into cohesive social groups (i.e., network fragmentation). Specifically, we find that social networks of relatively solitary species tend to demonstrate the highest degree heterogeneity, that social networks of gregarious species tend to be the most fragmented, and that socially hierarchical species are least clustered in their interactions. The structural differences between the social systems were detected after controlling for systematic biases in the data-collection (that might generate non-biological differences between the social structures). This suggests that the underlying differences in social network structures associated with each social system are biologically significant.

Social species are typically assumed to have a skewed degree distribution (for e.g. bottlenose dolphins Lusseau *et al.* (2003), wire-tailed manakins Ryder *et al.* (2008)), which implies that a small proportion of individuals have a large number of social partners. Our results, however, show that degree heterogeneity in relatively solitary species can be much higher than social species. Large variation in the number of social connections in relatively solitary species may simply arise due to a high variation in spatial behavior as compared to social species (Pinter-Wollman, 2015; Sah *et al.*, 2016). A homogeneous degree distribution in socially hierarchical species, such as ants and savanna baboons, could allow for efficient and equitable information transfer to all individuals (Blonder & Dornhaus, 2011; Cantor & Whitehead, 2013). Low average clustering coefficient, as observed in socially hierarchical species, indicates that an individual’s local social network is not tightly interconnected (i.e., individual’s contacts do not form a tight clique), and is known to increase network resilience and stability in response to perturbations such as the removal or death of individuals (Flack *et al.*, 2006; Krause *et al.*, 2014).

Our results also show that social networks of gregarious species are the most subdivided (but not disconnected) into cohesive social subgroups. The presence of many but small, socially cohesive subgroups within social networks of gregarious species can be explained based on the behavioural tendency to switch affiliative partners; as a result, individuals form consistent social bonds with a only small subset of individuals (Rubenstein *et al.*, 2015). Many gregarious species also form groups based on sex or age class, kinship and functional roles (Kanngiesser *et al.*, 2011) or due to high spatial or temporal variability in resources (Couzin, 2006; Couzin & Laidre, 2009; Sueur *et al.*, 2011). Previous theoretical models have shown that modular subdivisions promote behavioural diversity and cooperation (Whitehead & Lusseau, 2012; Gianetto & Heydari, 2015). Gregarious species may therefore limit the size of their social subgroups to maximize benefits of cooperation, making their social networks subdivided (Marcoux & Lusseau, 2013).

Our results show that the observed structural differences between the three social systems are primarily driven by the presence weak ties in their social networks. The reason why filtering out weak weighted edges removes most structural differences between social systems lies in their organization of weak ties. Individuals of social species disproportionately allocate effort among their social connections in order to maintain overall group connectivity (Fig. S1, Supporting information) and are also known to have high social fluidity (Colman & Bansal, 2017). Removing weak ties from networks of social species therefore increases variation in individual connectivity (degree heterogeneity), with a relatively minor decrease in their global connectivity (average betweenness centrality). Consequently, the global connectivity of social species in 10%-15% thresholded networks is significantly higher than relatively solitary species.

Previous studies have typically focused on group size as the key parameter impacting disease transmission and group living costs. However, the expectation of higher disease costs of group living has yielded mixed results (Arnold & Anja, 1993; Rifkin, Nunn & Garamszegi, 2012; Patterson & Ruckstuhl, 2013), which can be explained in part by the presence of group-level behavioural (Meunier, 2015; Schaller *et al.*, 2015) and physiological defense (Habig, Archie & Habig, 2015) against infection spread, as well as the presence of chronic social stress (Kappeler *et al.*, 2015; Nunn *et al.*, 2015). While group size might be easy parameter to measure, it does not capture the complex spatio-temporal dynamics of most animal societies. By performing disease simulations over empirical networks with different interaction types, we consider a range of infectious diseases with different transmission routes, including those that spread by direct contact, and those that spread by asynchronous contact between individuals in a population. Our analysis shows that the organization of social patterns may not provide general protection against pathogens of a range of transmission potential. We note that our results on epidemic size and duration are specific to pathogens that follow SIR (susceptible-infected-recovered) infection dynamics. The outcome of epidemic probability, however, is expected to be similar across different models of infectious disease spread (such as infections that provide temporary immunity or chronic infections).

We find that socially hierarchical species experience longer outbreaks of low transmissibility infections and frequent epidemics of moderately contagious infections because of low variation in individual and local connectivity (i.e., degree heterogeneity and average clustering coefficient) as compared to other social systems. Networks with low degree heterogeneity are known to experience steady protracted outbreaks, in contrast to explosive rapid outbreaks fueled by super-spreaders in high degree heterogeneity networks (Meyers *et al.*, 2005; Kiss, Green & Kao, 2006; Bansal, Grenfell & Meyers, 2007). High average clustering coefficient is also believed to create redundant paths between individuals making it harder for slow spreading infections to encounter new susceptible individuals and percolate throughout the network, prolonging infection spread (Newman, 2003).

In our disease simulations, highly fragmented social networks of gregarious species experienced frequent epidemics of highly contagious infections, and longer epidemics of moderately to highly transmissible pathogens. Our recent work has shown that infection spread in highly fragmented networks gets localized within socially cohesive subgroups (structural trapping), which enhances local transmission but causes structural delay of global infection spread (Sah *et al.*, 2017). In addition, our results suggest that highly transmissible pathogens are able to avoid stochastic extinction in fragmented networks by reaching "bridge" nodes, but experience delay in transmission due to the presence of structural bottlenecks.

As this study involved comparisons of social networks across a broad range of taxonomic groups and data sampling methods, we made a number of assumptions that could shape the results. First, because the impact of edge weights on disease transmission can be context-dependent, depending on the type of inter-action, transmission mode of pathogen, and the relative time scale of network collection and pathogen spread, we have chosen to not include edge weights while performing our computational disease experiments. Future meta-analytic studies can leverage a growing number of transmission studies to explicitly incorporate the role of contact intensity on disease transmission (Aiello *et al.*, 2016; Manlove *et al.*, 2017). Second, we assume that social contacts remain unaltered after an infection is introduced in population. Presence of infection, however, can alter the social connectivity of hosts (Croft *et al.*, 2011; Lopes, Block & König, 2016). Future species specific studies can take advantage of host specific experimental manipulations, where possible, to gain in-depth insight towards host behavior - infection feedback (Ezenwa *et al.*, 2016; Silk *et al.*, 2017a). Finally, in our network database there were some systematic differences in data-collection methodologies across social systems. Specifically, all data for relative solitary species were collected by sampling individuals over a specified spatial range, because definition of social groups for these species can be vague. As observations of direct interactions in relatively solitary species are rare, all networks of relatively solitary species in our database were based on direct or indirect spatial associations. Although the meta-analysis described in this study controlled for such biases in data-collection, the results should be interpreted as a conceptual understanding about the differences between the social systems in terms of empirical networks that have been published in the literature, and not as a general prediction about the differences in social systems.

Overall, our results suggests that the organization of social networks in gregarious species are more efficient in preventing outbreaks of moderately contagious pathogens than socially hierarchical species. Conversely, networks of socially hierarchical species experience fewer outbreaks of fast spreading infectious diseases as compared to gregarious species. The question of why this is so warrants detailed future investigations of the eco-evolutionary trajectory of social connectivity in the two social systems. It is likely that the organization of social networks in socially hierarchical species may have evolved to prevent outbreaks of highly transmissible pathogens, while relying on alternate group-level disease defense mechanisms (including sanitary behaviors, allogrooming, and the use of antimicrobials) to prevent outbreaks of low to moderate transmissibility infections. Since the social networks included in the meta-analysis were selected regardless of the presence of infectious diseases in the populations, the organization of network structure could also reflect the selection pressure of past infections, presence of other ecological/evolutionary drivers (Pinter-Wollman *et al.*, 2013), or conflicting selection pressures posed by the effort to maximize transmission of information.

## Challenges and opportunities

The sociality of animal species has been traditionally classified based on qualitative phenotypes and life history traits, and the classification typically differs between taxonomic groups. While this categorization scheme is convenient, it does not capture the continuum of social behaviour. As a step forward, recent studies have proposed quantitative indices of sociality (Silk, Altmann & Alberts, 2006; Aviles & Harwood, 2012). The results of our study support the potential use of network structure as a means of quantifying social complexity across taxonomic boundaries. Similar predictive meta-analyses can also be used to identify species that are outliers in the current sociality classification system based on the organization of their social structure.

However, we need to overcome several challenges before robust comparative analysis can be performed on social networks across broad taxonomic groups to address such issues. First, comparing network structure across taxonomic groups where data is aggregated over different spatio-temporal scales is challenging. Aggregating interactions over small time-periods may omit important transient inter-actions, whereas aggregating data over long time-periods may lead to a saturated network where distinguishing social organization may be difficult. Spatial constraints and environmental heterogeneity can also impose a considerable influence on the social network structure (Davis *et al.*, 2015; Leu *et al.*, 2016). Additionally, the consideration of relative time scale of animal interaction and infectious period of pathogen is critical in making accurate predictions of disease spread. Future comparative studies should therefore consider standardizing data over temporal and spatial scales.

The second challenge lies in effectively controlling for inherent biases in data collection methodologies across taxonomic groups. As direct observation of interactions is difficult in relatively solitary species, social networks are usually constructed based on direct or indirect spatial associations (rather than interactions) between individuals in a population (rather than a local group). Network size correlates to sampling intensity in many cases, and is therefore a poor proxy to group size. Social network studies of relatively solitary species are also relatively sparse compared to social species.

The third challenge for comparative studies of animal social networks is utilizing data-sources published in inconsistent formats. To facilitate in-depth meta-analyses of network data, we encourage researchers to accompany animal network datasets with the following details: data sampling method, location of the data collection, type of population monitored (captive, semi-captive, free-ranging), edge definition, edge weighting criteria, node attributes (such as demography), temporal resolution of data, temporal and spatial aggregation of the data, proportion of animals sampled in the area, and population density. When exact measurements of these data attributes are difficult, using reasonable approximations or proxies would be more useful than no information.

## Conclusions

In summary, our study broadens the scope of network analysis from being just species-specific to a meta-analytic approach, and provides new insights towards how the organization of interaction patterns can mediate disease costs of sociality. We note that there is enormous potential of adopting a comparative approach to study the commonalities and differences in social networks across a wide range of taxonomic groups and social systems. Future studies can use this approach to quantitatively test several evolutionary and ecological hypotheses, including ones on the tradeoffs of group living, the contributions of social complexity to intelligence, the propagation of social information, and social resilience to population stressors.

## Acknowledgements

We thank José David Méndez for his assistance in organizing data. We are grateful for the constructive and insightful reviews provided by Matthew Silk, Damien Farine and two anonymous reviewers; as well as for the feedback provided by Stephan Leu on a previous version of this manuscript. This work was supported by the National Science Foundation Ecology and Evolution of Infectious Diseases grant 1216054.

## Data accessibility

The data for all animal social network measures used in the study, and references where the actual network can be accessed, is available through the Bansal Lab Dataverse at (link).

## References

Aiello, C.M., Nussear, K.E., Esque, T.C., Emblidge, P.G., Sah, P., Bansal, S. & Hudson, P.J. (2016) Host contact and shedding patterns clarify variation in pathogen exposure and transmission in threatened tortoise *Gopherus agassizii*: implications for disease modeling and management. Journal of Animal Ecology, 85, 829–842. ISSN 00218790.

Altizer, S., Nunn, C.C.L., Thrall, P.P.H., Gittleman, J.L.J., Antonovics, J., Cunningham, A.A.a., Dobson, A.A.P.A., Ezenwa, V., Jones, K.K.E.K., Pedersen, A.A.B., Poss, M. & Pulliam, J.J.R.J. (2003) Social organization and parasite risk in mammals: Integrating theory and empirical studies. Annual Review of Ecology, Evolution, and Systematics, 34, 517–547.

Anderson, R.M., May, R.M. & Anderson, B. (1992) Infectious diseases of humans: dynamics and control, vol. 28. Wiley Online Library.

Arnold, W. & Anja, V.L. (1993) Ectoparasite loads decrease the fitness of alpine marmots (Marmota marmota) but are not a cost of sociality. Behavioral Ecology, 4, 36–39.

Aureli, F., Schaffner, C.M., Boesch, C., Bearder, S.K., Call, J., Chapman, C.A., Connor, R., Fiore, A.D., Dunbar, R.I.M., Henzi, S.P., Holekamp, K., Korstjens, A.H., Layton, R., Lee, P., Lehmann, J., Manson, J.H., Ramos-Fernandez, G., Strier, K.B. & Schaik, C.P.v. (2008) Fission-Fusion Dynamics. Current Anthropology, 49, 627–654.

Aviles, L. & Harwood, G. (2012) A Quantitative Index of Sociality and Its Application to Group-Living Spiders and Other Social Organisms. Ethology, 118, 1219–1229.

Bailey, N.T. (1957) The mathematical theory of epidemics.

Bansal, S., Grenfell, B.T. & Meyers, L.A. (2007) When individual behaviour matters: homogeneous and network models in epidemiology. Journal of the Royal Society, Interface / the Royal Society, 4, 879–91. ISSN 1742-5689.

Blondel, V.D., Guillaume, J.L., Lambiotte, R. & Lefebvre, E. (2008) Fast unfolding of communities in large networks. Journal of Statistical Mechanics: Theory and Experiment, 2008, P10008.

Blonder, B. & Dornhaus, A. (2011) Time-ordered networks reveal limitations to information flow in ant colonies. PloS one, 6, e20298. ISSN 1932-6203.

Cantor, M. & Whitehead, H. (2013) The interplay between social networks and culture: theoretically and among whales and dolphins The interplay between social networks and culture: theoretically and among whales and dolphins. Philosophical transactions of the Royal Society of London. Series B, Biological sciences, 368, 1–10. ISSN 0962-8436, 1471-2970.

Colman, E. & Bansal, S. (2017) Social fluidity mobilizes infectious disease in human and animal populations. bioRxiv.

Couzin, I.D. & Laidre, M.E. (2009) Fission-fusion populations. Current Biology, 19, 633–635. ISSN 09609822.

Couzin, I. (2006) Behavioral ecology: social organization in fission–fusion societies. Current Biology, 16, 169–171. ISSN 0960-9822. URL http://www.sciencedirect.com/science/article/pii/S0960982206011924

Craft, M.E. (2015) Infectious disease transmission and contact networks in wildlife and livestock. Philosophical Transactions of the Royal Society of London. Series B, Biological Xciences, 370, 1–12. ISSN 1471-2970.

Cremer, S., Armitage, S.a.O. & Schmid-Hempel, P. (2007) Social immunity. Current biology: CB, 17, R693–702.

Croft, D.P., Edenbrow, M., Darden, S.K., Ramnarine, I.W., Oosterhout, C.v. & Cable, J. (2011) Effect of gyrodactylid ectoparasites on host behaviour and social network structure in guppies Poecilia reticulata. Behavioral Ecology and Sociobiology, 65, 23–35.

Croft, D.P., James, R. & Krause, J. (2008) Exploring Animal Social Networks. Princeton University Press.

Davis, S., Abbasi, B., Shah, S., Telfer, S. & Begon, M. (2015) Spatial analyses of wildlife contact networks. Journal of the Royal Society, Interface, 12.

Ezenwa, V.O., Ghai, R.R., McKay, A.F. & Williams, A.E. (2016) Group living and pathogen infection revisited. Current Opinion in Behavioral Sciences, 12, 66–72. ISSN 23521546.

Farine, D.R. & Whitehead, H. (2015) Constructing, conducting, and interpreting animal social network analysis. The Journal of animal ecology, pp. 1144–1163.

Faust, K. (2006) Comparing Social Networks: Size, Density, and Local Structure. Metodološki zvezki, 3, 185–216.

Faust, K. & Skvoretz, J. (2002) Comparing Networks across Space and Time, Size and Species. Sociological Methodology, 32, 267–299. ISSN 0081-1750.

Flack, J.C., Girvan, M., de Waal, F.B.M. & Krakauer, D.C. (2006) Policing stabilizes construction of social niches in primates. Nature, 439, 426–429. ISSN 0028-0836.

Gelman, A. & Rubin, D.B. (1992) Inference from iterative simulation using multiple sequences. Statistical science, pp. 457–472.

Gelman, A. (2006) Prior distributions for variance parameters in hierarchical models (Comment on Article by Browne and Draper). Bayesian Analysis, 1, 515–534.

Gianetto, D.A. & Heydari, B. (2015) Network Modularity is essential for evolution of cooperation under uncertainty. Scientific Reports, 5, 9340.

Godfrey, S.S., Bull, C.M., James, R. & Murray, K. (2009) Network structure and parasite transmission in a group living lizard, the gidgee skink, Egernia stokesii. Behavioral Ecology and Sociobiology, 63, 1045–1056.

Grenfell, B.T. & Dobson, A.P. (1995) Ecology of infectious diseases in natural populations, vol. 7. Cambridge University Press.

Habig, B., Archie, E.A. & Habig, B. (2015) Social status, immune response and parasitism in males: a meta-analysis. Philosophical Transactions of the Royal Society B, 370, 20140109. ISSN 1471-2970.

Hadfield, J. (2014) MCMCglmm course notes.

Hadfield, J.D. (2010) MCMC methods for multi-respoinse generalized linear mixed models: The MCMCglmm R package. Journal of Statistical Software, 33, 1–22.

Heffernan, J., Smith, R. & Wahl, L. (2005) Perspectives on the basic reproductive ratio. Journal of The Royal Society Interface, 2, 281–293. ISSN 1742-5689.

Hock, K. & Fefferman, N.H. (2012) Social organization patterns can lower disease risk without associated disease avoidance or immunity. Ecological Complexity, 12, 34–42.

Kanngiesser, P., Sueur, C., Riedl, K., Grossmann, J. & Call, J. (2011) Grooming network cohesion and the role of individuals in a captive chimpanzee group. American Journal of Primatology, 73, 758–767.

Kappeler, P.M. & van Schaik, C.P. (2002) Evolution of Primate Social Systems. International Journal of Primatology, 23, 707–740. ISSN 1573-8604.

Kappeler, P.M., Cremer, S., Nunn, C.L. & Kappeler, P.M. (2015) Sociality and health: impacts of sociality on disease susceptibility and transmission in animal and human societies. Philosophical transactions of the Royal Society of London. Series B, Biological sciences, 370, 20140116. ISSN 0962-8436.

Keeling, M. (2005) The implications of network structure for epidemic dynamics. Theoretical population biology, 67, 1–8. ISSN 0040-5809.

Kiss, I.Z., Green, D.M. & Kao, R.R. (2006) Infectious disease control using contact tracing in random and scale-free networks. Journal of the Royal Society, Interface / the Royal Society, 3, 55–62.

Kiss, I.Z., Miller, J.C. & Simon, P.L. (2017) Mathematics of epidemics on networks: from exact to approximate models, vol. 46. Springer.

Krause, J., Croft, D.P. & James, R. (2007) Social network theory in the behavioural sciences: Potential applications. Behavioral Ecology and Sociobiology, 62, 15–27. ISSN 03405443.

Krause, J., James, R., Franks, D.W. & Croft, D.P., eds. (2014) Animal social networks. Oxford University Press, USA.

Lenth, R.V. (2016) Least-squares means: the R package lsmeans. Journal of Statistical Software, 69, 1–33.

Leu, S.T., Farine, D.R., Wey, T.W., Sih, A. & Bull, C.M. (2016) Environment modulates population social structure: experimental evidence from replicated social networks of wild lizards. Animal Behaviour, 111, 23–31.

Leu, S.T., Kappeler, P.M. & Bull, C.M. (2010) Refuge sharing network predicts ectoparasite load in a lizard. Behavioral Ecology and Sociobiology, 64, 1495–1503.

Loehle, C. (1995) Social Barriers to Pathogen Transmission in Wild Animal Popluations. Ecology, 76, 326–335.

Lopes, P.C., Block, P. & König, B. (2016) Infection-induced behavioural changes reduce connectivity and the potential for disease spread in wild mice contact networks. Scientific Reports, 6, 31790. ISSN 2045-2322.

Lusseau, D., Schneider, K., Boisseau, O.O.J., Haase, P., Slooten, E. & Dawson, S.S.M. (2003) The bottlenose dolphin community of Doubtful Sound features a large proportion of long-lasting associations. Behavioral Ecology and Sociobiology, 54, 396–405.

Manlove, K.R., Cassirer, E.F., Plowright, R.K., Cross, P.C. & Hudson, P.J. (2017) Contact and contagion: Probability of transmission given contact varies with demographic state in bighorn sheep. Journal of Animal Ecology, 86, 908–920. ISSN 13652656.

Marcoux, M. & Lusseau, D. (2013) Network modularity promotes cooperation. Journal of Theoretical Biology, 324, 103–108.

Meunier, J. (2015) Social immunity and the evolution of group living in insects. Philosophical Transactions B of the Royal Society, 370, 20140102.

Meyers, L.A., Pourbohloul, B., Newman, M.E.J., Skowronski, D.M. & Brunham, R.C. (2005) Network theory and SARS: predicting outbreak diversity. Journal of theoretical biology, 232, 71–81.

Newman, M. (2003) Properties of highly clustered networks. Physical Review E, 68, 26121.

Nunn, C.L., Craft, M.E., Gillespie, T.R., Schaller, M., Kappeler, P.M. & Nunn, C.L. (2015) The sociality – health – fitness nexus: synthesis, conclusions and future directions. Philosophical Transactions of the Royal Society B, 370, 20140115. ISSN 1471-2970.

Otterstatter, M.C. & Thomson, J.D. (2007) Contact networks and transmission of an intestinal pathogen in bumble bee (Bombus impatiens) colonies. Oecologia, 154, 411–21.

Patterson, J.E.H. & Ruckstuhl, K.E. (2013) Parasite infection and host group size: a meta-analytical review. Parasitology, pp. 1–11.

Pinter-Wollman, N. (2015) Persistent variation in spatial behavior affects the structure and function of interaction networks. Current Zoology, 61, 98–106.

Pinter-Wollman, N., Hobson, E.A., Smith, J.E., Edelman, A.J., Shizuka, D., de Silva, S., Waters, J.S., Prager, S.D., Sasaki, T., Wittemyer, G., Fewell, J. & McDonald, D.B. (2013) The dynamics of animal social networks: analytical, conceptual, and theoretical advances. Behavioral Ecology, 25, 242–255. ISSN 1045-2249.

Plummer, M., Best, N., Cowles, K. & Vines, K. (2006) CODA: Convergence diagnosis and output analysis for MCMC. R news, 6, 7–11.

Prange, S., Gehrt, S.D., Hauver, S. & Voigt, C.C. (2011) Frequency and duration of contacts between free-ranging raccoons: uncovering a hidden social system. Journal of Mammalogy, 92, 1331–1342.

Rifkin, J.L., Nunn, C.L. & Garamszegi, L.Z. (2012) Do Animals Living in Larger Groups Experience Greater Parasitism? A Meta-Analysis. The American Naturalist, 180, 70–82.

Rubenstein, D.I., Sundaresan, S.R., Fischhoff, I.R., Tantipathananandh, C. & Berger-wolf, T.Y. (2015) Similar but Different: Dynamic Social Network Analysis Highlights Fundamental Differences between the Fission-Fusion Societies of Two Equid Species, the Onager and Grevy ' s Zebra. PLoS ONE, 10, 1–21.

Ryder, T.B., McDonald, D.B., Blake, J.G., Parker, P.G. & Loiselle, B.A. (2008) Social networks in the lek-mating wire-tailed manakin (Pipra filicauda). Proceedings of the Royal Society B-Biological Sciences, 275, 1367–1374.

Sah, P., Leu, S.T., Cross, P.C., Hudson, P.J. & Bansal, S. (2017) Unraveling the disease consequences and mechanisms of modular structure in animal social networks. Proceedings of the National Academy of Sciences of the United States of America, 114, 4165–4170. ISSN 1091-6490.

Sah, P., Nussear, K.E., Esque, T.C., Aiello, C.M., Hudson, P.J. & Bansal, S. (2016) Inferring social structure and its drivers from refuge use in the desert tortoise, a relatively solitary species. Behavioral Ecology and Sociobiology, pp. 1–13.

Sah, P., Singh, L.O., Clauset, A. & Bansal, S. (2014) Exploring community structure in biological networks with random graphs. BMC bioinformatics, 15, 220.

Schaller, M., Murray, D.R., Bangerter, A. & Schaller, M. (2015) Implications of the behavioural immune system for social behaviour and human health in the modern world. Philosophical Transactions of the Royal Society B, 370, 20140105. ISSN 1471-2970.

Schielzeth, H. (2010) Simple means to improve the interpretability of regression coeffcients. Methods in Ecology and Evolution, 1, 103–113.

Silk, J., Cheney, D. & Seyfarth, R. (2013) A practical guide to the study of social relationships. Evolutionary Anthropology, 22, 213–225.

Silk, J.B., Altmann, J. & Alberts, S.C. (2006) Social relationships among adult female baboons (papio cynocephalus) I. Variation in the strength of social bonds. Behavioral Ecology and Sociobiology, 61, 183–195.

Silk, M.J., Croft, D.P., Delahay, R.J., Hodgson, D.J., Boots, M., Weber, N. & McDonald, R.A. (2017a) Using Social Network Measures in Wildlife Disease Ecology, Epidemiology, and Management. BioScience, 67, 245–257. ISSN 0006-3568.

Silk, M.J., Croft, D.P., Delahay, R.J., Hodgson, D.J., Weber, N., Boots, M. & McDonald, R.A. (2017b) The application of statistical network models in disease research. Methods in Ecology and Evolution. ISSN 2041210X.

Silk, M.J., Croft, D.P., Tregenza, T. & Bearhop, S. (2014) The importance of fission – fusion social group dynamics in birds. Ibis, 156, 701–715. ISSN 00191019.

Slater, P.J.B. & Halliday, T.R., eds. (1994) Behavior and Evolution. Cambridge University Press, USA.

Stroeymeyt, N., Casillas-Pérez, B. & Cremer, S. (2014) Organisational immunity in social insects. Current Opinion in Insect Science, 5, 1–15. ISSN 22145745.

Sueur, C., King, A.J., Conradt, L., Kerth, G., Lusseau, D., Mettke-Hofmann, C., Schaffner, C.M., Williams, L., Zinner, D. & Aureli, F. (2011) Collective decision-making and fission-fusion dynamics: A conceptual framework. Oikos, 120, 1608–1617. ISSN 00301299.

VanderWaal, K.L. & Ezenwa, V.O. (2016) Heterogeneity in pathogen transmission: mechanisms and methodology. Functional Ecology, pp. n/a–n/a. ISSN 02698463.

White, L.A., Forester, J.D. & Craft, M.E. (2015) Using contact networks to explore mechanisms of parasite transmission in wildlife. Biological Reviews. ISSN 14647931.

Whitehead, H. & Lusseau, D. (2012) Animal social networks as substrate for cultural behavioural diversity. Journal of Theoretical Biology, 294, 19–28.

